# Memory T cell rapid recall is driven by memory-specific AP-1 recruitment determined by epigenome and co-factor interactions

**DOI:** 10.64898/2026.02.13.705382

**Authors:** Adenike R. Shittu, Andrew VonHandorf, Michael Kotliar, Valerii Pavlov, Sarah Potter, Xiaoting Chen, Mathew T. Weirauch, Artem Barski

## Abstract

CD4 T cell memory is essential for long-lasting protective immunity to repeat infections. Unlike naïve T cells, memory cells possess rapid recall ability to quickly produce effector molecules in response to antigen re-exposure. This ability was shown to be associated with epigenetic gene poising. Here, we examine how the activation-inducible transcription factors, AP□1 and NF□κB, regulate rapid recall gene expression. We found that AP-1 is required for their induction and that the enhanced induction of rapid recall genes in memory cells is associated with memory-specific binding of AP□1. Memory-specific AP□1 binding, in turn, is enabled by enhanced chromatin accessibility and reduced DNA methylation at regulatory elements. As the AP-1 DNA-binding motif itself does not contain methylatable CpGs, methylation likely affects the binding of AP-1 co-factors, such as ETS proteins, or accessibility of the region in general. Finally, both common and memory-specific AP□1/NF□κB binding sites show strong overlap with autoimmune and inflammatory disease risk variants, highlighting the clinical relevance of memory T cell epigenetic regulation.

## INTRODUCTION

Memory T cells provide long-term immune protection by rapidly mounting a robust response upon re-exposure to their cognate antigens^1^. Memory T cells are broadly categorized into effector memory T cells (TEM) and central memory T cells (TCM), which differ in their proliferative capacity, tissue localization, and effector capability^2^. Both subsets collectively sustain long-term immune surveillance and protection. Memory T cells rapidly produce cytokines and effector molecules upon restimulation, thus enabling a protective response^3^. The molecular mechanism that underlies the enhanced response in memory T cells compared to naive T cells is not fully understood^4^. Potentially, epigenetic modifications, T cell receptor (TCR) signaling pathways, transcription factors (TFs) activities, metabolic, and post-transcriptional mechanisms may all contribute to rapid recall immune responses in memory T cells^5^.

Upon recognition of cognate antigen via engagement of the T cell receptor (TCR) and associated costimulatory receptors, naive and memory T cells become activated, triggering a cascade of intracellular signaling events, chromatin remodeling, and activation of transcription. Compared to naive T cells, memory T cells exhibit reduced levels of tyrosine-phosphorylated signaling intermediates and increased levels of phosphatases following T cell activation, leading to decreased activation signal and, thus, suggesting that the enhanced immune response cannot be explained by increased signaling^6–9^. Downstream, memory T cells exhibit reduced DNA methylation at the loci of immune response related genes^10,11^, more open chromatin configuration, and permissive histone modifications that are associated with rapid transcriptional activation upon antigen re-encounter^12^. Collectively, these findings highlight distinct phenotypic and molecular differences in the epigenetic landscapes between naive and memory T cells. However, the molecular mechanisms linking these epigenetic alterations to the rapid recall capacity of memory T cells remain elusive.

AP-1 and NF-κB are T cell activation-inducible TFs that translocate to the nucleus upon activation and induce the expression of cytokines, effector molecules, and other immune response genes^13–17^. Naive T cells undergo extensive chromatin remodeling upon activation. AP-1 is required for this chromatin remodeling at the majority of opening sites^18^. However, memory T cells already possess pre-accessible chromatin at many of these loci, supposedly allowing immediate gene activation^12,19^. Although the activity of AP-1 is well-characterized in naive T cells, its specific roles in shaping chromatin architecture and gene expression in memory subsets remain less defined. We hypothesize that differential binding of activation-inducible TFs, such as AP-1, influences epigenetic modifications and drives the rapid recall response in memory T cells. Identifying key TF regulators and mechanisms of AP-1 activity in rapid recall response in memory T cells can guide development of strategies to target memory cells in vivo that could improve vaccine design, boost immune defenses in infection, and guide CAR T cell therapies in autoimmune, allergic diseases, and cancers.

Herein, we studied the role of AP-1 and NF κB in the activation-induced transcription and chromatin remodeling that underlie rapid recall responses in TEM and TCM cells. To functionally inhibit AP-1 activity, we employed a dominant-negative protein, A-FOS^20^, which broadly prevents the AP-1 heterodimer formation and its DNA-binding activity of the AP-1 TFs. We performed bulk RNA-seq and ATAC-seq to assess how AP-1 inhibition alters gene expression and chromatin accessibility. In addition, we conducted ChIP-seq to evaluate the binding of AP-1 (FRA2, JUNB subunits) and NF κB (*NFKB1*, p50) in memory cells compared to naive. We show that AP-1 binding is required for the induction of rapid-recall genes in memory T cells. Further, we show that stronger inducibility of rapid recall genes in memory T cells is associated with memory-specific AP-1 differential binding. Differential binding, in turn, is driven by differential DNA accessibility and methylation around rapid recall genes. Together, these findings demonstrate that AP-1 and NF-κB are critical regulators of the transcriptional program that enable rapid recall responses in memory CD4 T cells. Notably AP-1 plays a dual role in transcriptional activation and chromatin remodeling, highlighting its potential as a therapeutic target for the development of next-generation vaccines and treatments for infectious and autoimmune diseases.

## RESULTS

### Experimental system to investigate the epigenetic and transcriptional activity of AP-1 in rapid recall response in memory T cells

To test the differential role AP-1 plays in CD4 T cell subset activation, human naive (CD4+ CD27+ CD45RO-), central (CD4+ CD27+ CD45RO+) and effector (CD4+ CD27- CD45RO+) memory CD4 T cells were isolated from the peripheral blood of healthy donors by negative magnetic enrichment for CD4 cells followed by flow sorting. Given that the human genome contains 18 AP-1 factors, the majority of which are induced upon T cell activation, we used a dominant-negative approach to inhibit AP-1 activity: recombinant AP-1 dominant-negative protein (A-FOS) was electroporated into resting cells prior to anti-CD3/28 bead stimulation (Fig. 1A, 1B). GFP protein was used as an electroporation control. A-FOS can bind and sequester JUN paralogs, preventing FOS–JUN complex formation^20–24^. Prior studies demonstrate that electroporated AFOS and GFP protein remain in CD4 T cells for >5 h^18,20^. The cells were then rested in the media for 2 h and then were either activated with anti-CD3/28 beads for 5 h or kept in the media for an additional 5 h. Gene expression was measured by bulk RNA-seq and chromatin accessibility by bulk ATAC-seq.

**Fig. 1:**
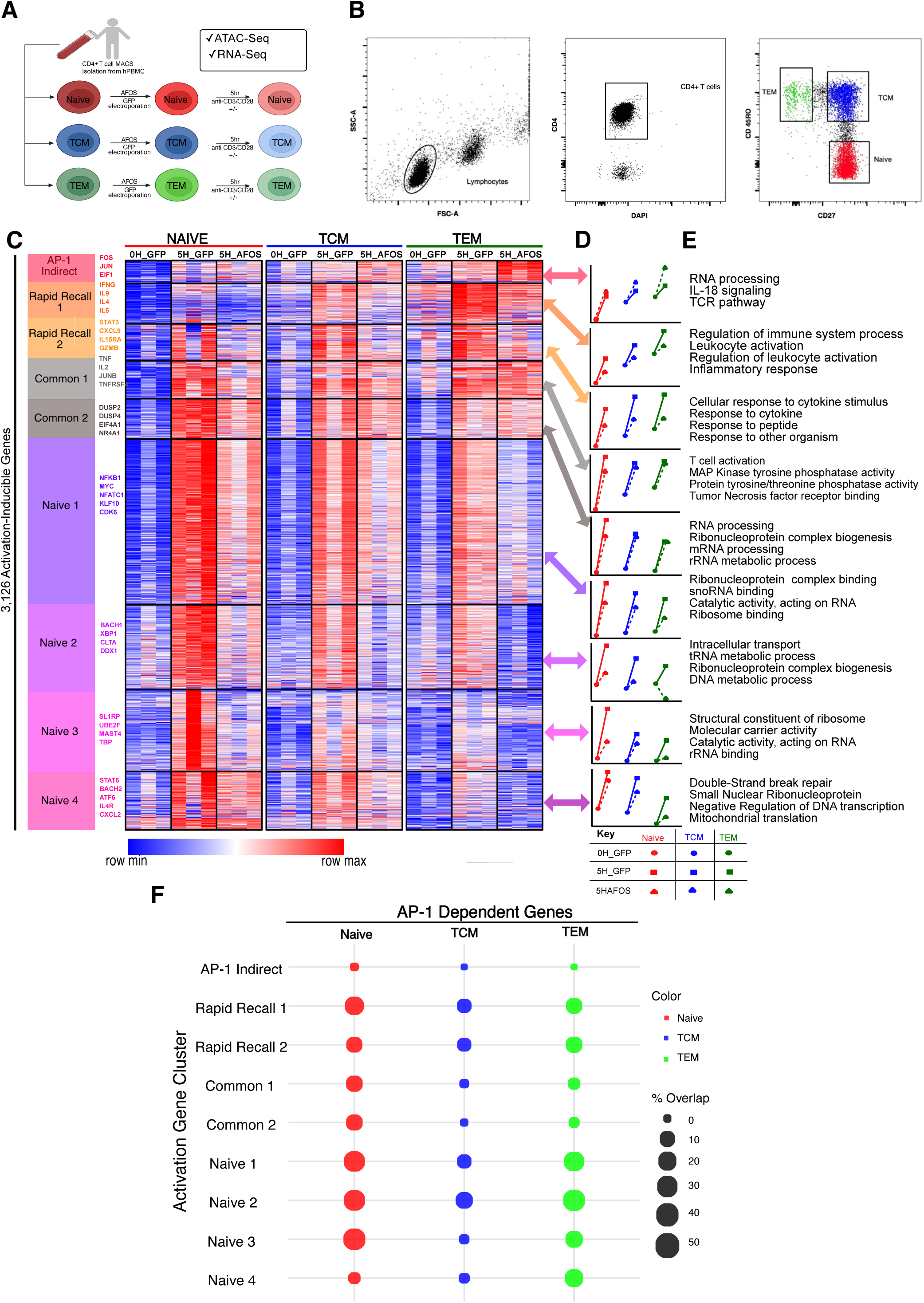
Inhibition of AP-1 alters the transcriptional program of activation-inducible and rapid recall genes in memory T cells. (**A.**) Experimental system used to investigate the AP-1‒dependent transcriptional program underlying rapid recall responses in human effector (TEM) and central (TCM) memory CD4 T cells. Sorted T cells were electroporated with GFP (control) or a dominant-negative A-FOS protein and either left unstimulated or stimulated with anti-CD3/CD28 activation beads for 5 h. (**B.**) Flow cytometry gating strategy for isolating naive, effector, and central memory CD4 T cells. (**C.**) Heatmap depicting mRNA expression levels of genes upregulated after 5-h activation in naive, effector, or central memory CD4 T cells. Genes were clustered using the expression patterns across naive and memory subsets as described in Methods. Minimum-maximum‒scaled, DESeq2-normalized expression counts are shown. (**D.**) Line plot showing Z score‒scaled average expression counts for each cluster across cell types and conditions. (**E.**) Gene ontology analysis of activation inducible gene clusters. **(F.**) Dot plot shows percentage of genes in each cluster from C whose expression is significantly changed (FDR<0.1; LFC>1.5) by A-FOS.

### Transcriptional changes in activated CD4 T cells are due to AP-1□dependent and □independent mechanisms in memory T cells

To understand CD4 T cell subset transcriptional changes, we compared activation-driven changes in three CD4 subsets (naive, TEM, and TCM) using bulk RNA-seq. Differential gene expression analysis revealed a widespread transcriptional change in naive, TEM, and TCM cell populations following the 5-h anti-CD3/CD28 stimulation. We identified 6,697 genes differentially expressed between resting and activated T cells across the 3 cell subtypes (|log_2_ fold change [LFC]| > 0.585, false discovery rate [FDR] < 0.1, DEseq2 Wald test^25^). Of these, 3,126 genes were upregulated in at least one cell type (Fig. 1C,1D), and 3,571 genes were downregulated upon activation (Supplementary Fig. S1A). We utilized clustering to further group activation inducible genes by subset-specific expression and the effect of AP-1 inhibition. Broadly, we identified clusters of rapid recall genes (r*apid recall 1 and 2*), which are highly induced in memory compared to naive T cells and include genes enriched in the inflammatory response and cytokines. Naive-specific genes with higher expression in activated naive cells (*naive 1, 2, 3, and 4)* are enriched in basic processes, including mRNA, tRNA, rRNA processing, and ribosomal complex biogenesis. Common genes (438 genes, induced at similar levels across naive and memory cells post stimulation; *common 1 and 2*) are enriched in T cell activation, RNA processing, and ribonucleoprotein complex biogenesis (Fig. 1C, 1D, 1E). We also identified an *AP-1 indirect* cluster of genes that were induced by activation in naive cells and stayed highly expressed in resting and activated memory cells. AP-1 inhibition did not suppress their induction in naive cells and enhanced it further in memory cells. This group included genes involved in RNA processing and the TCR pathway (Fig. 1C, 1D, 1E).

We further investigated genes undergoing transcriptional silencing during T cell activation and also categorized them into clusters (Supplementary Fig. S1A). The activation-repressed (c*ommon*) cluster comprises genes uniformly downregulated following stimulation across all cell types, indicative of a transcriptional silencing program during activation. Gene Ontology (GO) analysis revealed gene enrichment for DNA damage response, DNA repair, and protein ubiquitination (Supplementary Fig. S1B). The naive-repressed clusters (*naive*) include genes with high levels of expression in naive cells that become downregulated upon activation and are enriched for genes involved in intracellular transport (Supplementary Fig. S1A, 1B). Additionally, the *stable* cluster contains genes that maintain relative expression levels in memory cells upon activation, reflecting a preserved transcriptional state. Finally, *AP-1 indirect 1 and 2* cluster, comprised of genes that become downregulated upon activation but showed higher expression with the loss of AP-1, are enriched in pathways relevant to T cell biology, including the JNK signaling cascade, autophagy, and xenophagy (Supplementary Fig. S1A, 1B).

Next, we evaluated whether the expression of activation-induced genes was dependent on AP-1 by comparison of stimulated cells preincubated with the dominant negative A-FOS or GFP control cells. Upon 5 h of activation, differential gene expression analysis revealed a widespread transcriptional profile of 1,928 AP-1 dependent genes (|log_2_ FC| > 0.585, FDR < 0.1, DEseq2 Wald test) across the three CD4 T cell subtypes (Supplementary Fig. S2A). Broadly, inhibiting AP-1 by the A-FOS protein suppressed the expression of the activation-inducible genes, in particular those in the naive and rapid recall clusters (Fig. 1C, 1D, 1E, 1F and Supplementary Fig. 2A). Interestingly, induction of the genes in the common clusters (*common 1 and 2*) required AP-1 in naive cells but was no longer dependent on AP-1 in memory cells, as we observed little to no reduction in expression levels of *common 1 and 2* genes in memory cells with pretreatment with A-FOS (Fig. 1D, 1F). This suggested that AP-1 driven epigenetic reprograming during primary activation in the previous naive cell state was sufficient for their inducibility and transcription in memory T cells.

### The differential binding of AP-1 in memory cells regulates rapid recall response in memory T cells

We showed that the induction of rapid recall genes (*rapid recall 1 and 2*) in memory cells is dependent on AP-1 (Fig. 1C). However, AP-1 is present in the nuclei of both naive and memory cells upon activation. To explain the differential inducibility of rapid recall genes, we hypothesized that transcription of rapid recall genes in memory cells is due, at least in part, to differential binding of AP-1 at regulatory elements of rapid recall genes. To test this, we performed genome-wide profiling of DNA binding by ChIP-seq for two AP-1 family members, JUNB and FRA2, and another activation-inducible TF, NF κB (*NFKB1*, p50), in naive, TCM, and TEM cells after 5 h of anti-CD3/CD28 stimulation. We observed differential binding of JUNB, FRA2, and NF κB at the promoter and multiple enhancer regions of rapid recall genes, including cytokines, between the naive and memory subsets. For example, ChIP-seq showed binding of JUNB, FRA2, and NF κB at the promoter and multiple enhancer regions (5 kb upstream, 40 kb downstream, and 20 kb downstream) of *IFNG* in memory cells with little to no binding in naive cells (Fig. 2A).

**Fig. 2:**
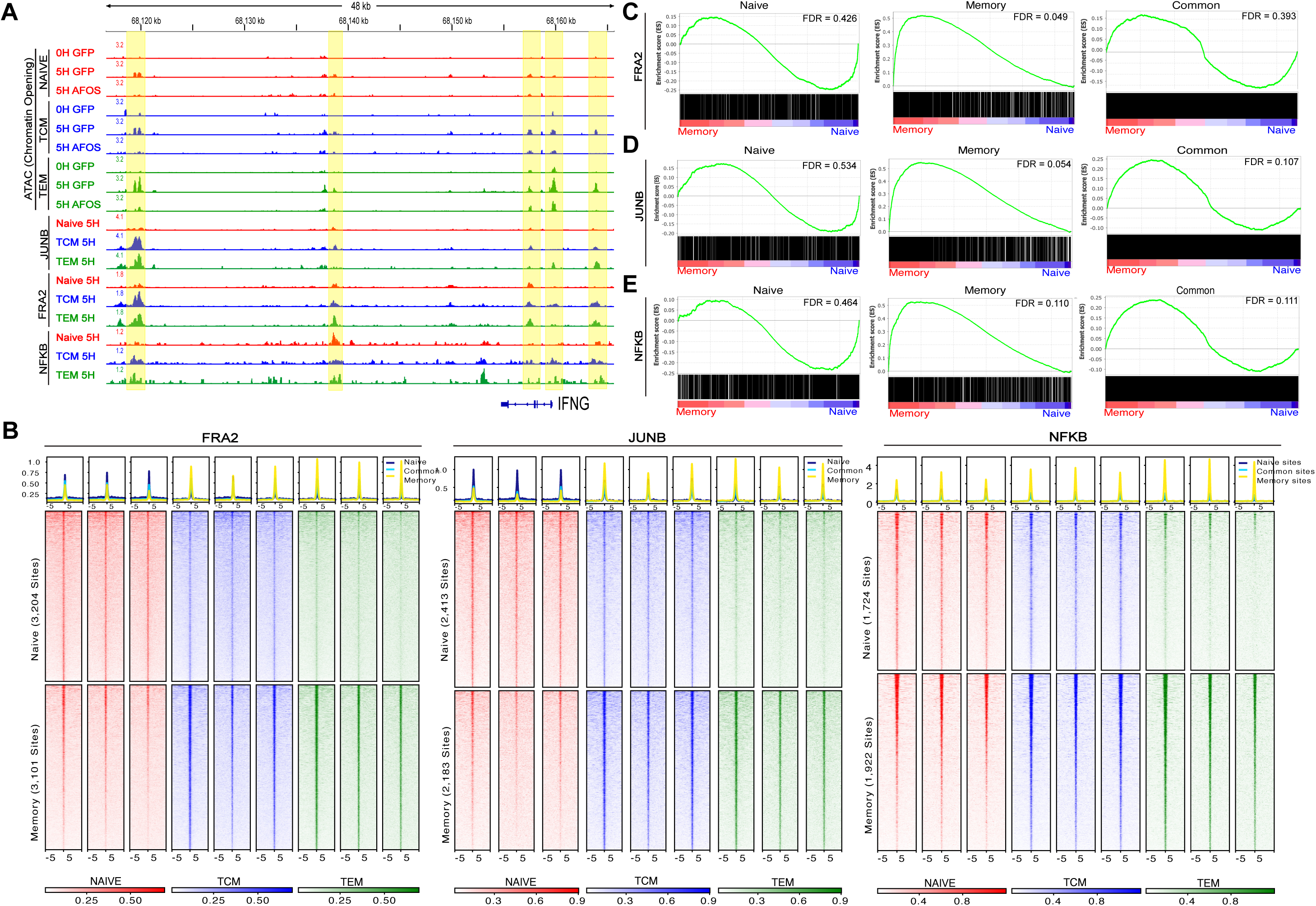
Differential binding of AP-1, NFAT, and NFKB in human memory CD4 T cells. (**A.**) Genome browser tracks showing ATAC-seq chromatin accessibility and binding of AP-1 (JUNB and FRA2) and NF-κB at the *IFNG* locus in naive, TEM, and TCM cells. Elements of interest are highlighted. (**B.**) Tag density plot illustrating the naive-specific and memory-specific binding sites of JUNB, FRA2, and NF-κB across naïve, TCM and TEM cell subsets. (**C-E.**) Gene set enrichment analysis shows that memory-specific JUNB-, FRA2-, and NF-κB‒bound regions are associated with higher expression in activated TEM than in naive cells. Transcription factor binding sites were assigned to genes by both distance and 3D structure as described in Methods. A ranked gene list was created by comparing gene expression in 5-h‒activated TEM vs. naive cells.

Next, we performed genome-wide differential binding analysis of JUNB, FRA2, and NF-κB and identified binding sites specific for naive and memory T cells (Fig. 2B) (|log_2_ FC| > 0.585, FDR < 0.1 DESeq2 Wald test within modified DiffBind) and binding sites common to all 3 cell populations (*common*, Supplementary Fig. S3). The majority (>80%) of the binding sites showed no difference between populations, suggesting that the role of AP-1 and NF κB is largely conserved across CD4 T cell subsets (see *EGR1*, *ETF1,* and *IL2RA* loci, Supplementary Fig. 4A, 4B). We identified 2,183 JUNB, 3,101 FRA2, and 1922 NF-κB sites that are memory-specific (enriched in TCM and TEM as compared to naive cells). Further, JUNB, FRA2, and NF κB ChIP-seq peaks were assigned to genes based on proximity (10 kb upstream and downstream) or 3D relationships derived from trac-loop^26^ and micro-C data^27^. We found that the genes associated with the shared memory-specific regions are enriched for cytokine production and immune responses to external stimuli (Supplementary Fig. S5C, S5F, S5I). Conversely, we extracted 2,413 JUNB, 3,204 FRA2, and 1,724 NF-κB sites that are bound in naive T cells (Fig. 2B). Pathway enrichment analysis revealed that these naive-specific regions are functionally associated with regulation of T cell activation, differentiation, and cell-cell adhesion, which are essential for early naive cell activation and migration in response to stimulation (Supplementary Fig. S5A, S5D, S5G). The common sites are enriched for protein transport, localization, and RNA biosynthesis (Supplementary Fig. S5B, S5E, S5H).

We further evaluated whether memory-specific AP-1 binding events are associated with the rapid recall transcriptional program in memory T cells. Gene set enrichment analysis^28^. showed that the genes associated with memory-specific binding of JUNB and FRA2 were significantly associated with higher expression in activated memory (TEM) than in naive cells. Association with differential NF-κB binding was trending but not statistically significant. Interestingly, naive-specific gene expression was not significantly associated with the naive-specific binding of these TFs (Fig. 2C, 2D, 2E).

### AP-1 regulates chromatin accessibility in memory CD4 T cells upon stimulation, influencing epigenetic reprogramming of regulatory regions of rapid recall genes

Having established that the induction of rapid recall genes in memory cells requires AP-1 activity and that enhanced inducibility of rapid recall genes is associated with memory-specific binding of AP-1, we next asked what drives AP-1 differential binding between naive and memory T cells. Differential binding can be regulated epigenetically either by chromatin accessibility or DNA methylation. Previously, we showed that AP-1 itself regulates chromatin accessibility of activation-inducible genes in naive cells^18^. Therefore, we expect that AP-1 regulates activation-driven chromatin opening in memory T cells as well. To test this, we utilized ATAC-seq to profile chromatin accessibility in naive, TCM, and TEM cells upon activation in the presence and absence of A-FOS; our analysis identified 14,357 regions with increased accessibility in at least one cell type and 874 regions with decreased accessibility (Fig. 3A, Supplementary Fig. S5A).

**Fig. 3:**
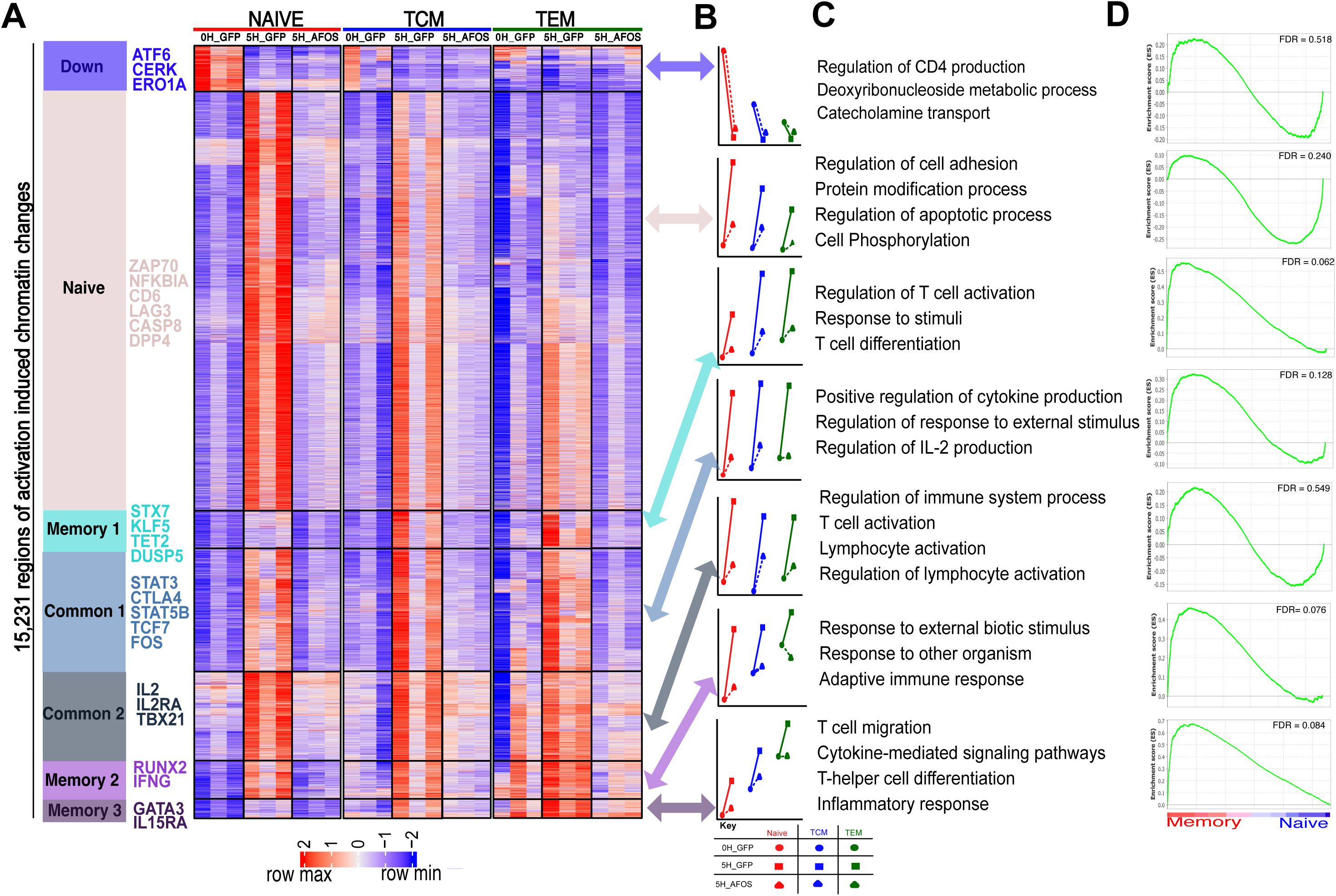
AP-1 influence chromatin accessibility of rapid recall genes in memory T cells. (**A.**) Heatmap showing genomic regions that gained or lost chromatin accessibility after 5-h stimulation in naive, TEM, and TCM cells. Regions are clustered using the accessibility patterns across naive and memory subsets. (**B.**) Line plot depicting Z score‒scaled average accessibility values for each cluster across cell types and conditions. (**C.**) Gene ontology analysis for each cluster of accessible regions. Peaks were assigned to genes as described in Methods. (**D.**) Gene set enrichment analysis shows association between the clusters and increased expression in activated memory cells as in Fig. 2E.

We next performed clustering of differentially accessible regions and identified regions with increased accessibility in either naive or memory activated cells. Of interest, regions in clusters *memory 2 and 3* were already more accessible in resting memory than naive cells and were opened further upon activation (Fig. 3A). *Memory 1* cluster was not accessible in resting memory cells and selectively opened only in memory cells, but not naive cells, upon activation. Gene set enrichment analysis (GSEA) of the nearest genes within *memory 1,2, and 3* clusters revealed that these regions are associated with memory-specific gene expression (Fig. 3D). In contrast, regions classified as *naive* were more accessible in naïve cells, and *common 1 and 2* regions showed similar accessibility induction across both naive and memory subsets. However, genes associated with these naive and common regions were not enriched for rapid recall or memory-specific programs. In all clusters with increased accessibility, AP-1 activity was required for chromatin remodeling.

### Poised chromatin and reduced DNA methylation in memory T cells influence the differential binding of AP-1 in memory T cells

Naive and memory CD4 T cells are well known to exhibit distinct epigenetic landscapes. Previous studies, including from our lab, have established that memory CD4 T cells have reduced DNA methylation and a chromatin state poised for immune response genes compared to those of naive T cells^10–12^. To determine whether the differential AP-1 and NF κB binding observed in memory T cells is attributable to preexisting open chromatin or reduced DNA methylation in memory T cells, we evaluated TF binding at the accessible regions described in Fig. 3A. We initially hypothesized that AP-1 binding in memory T cells would occur predominantly at regions with preestablished open or poised chromatin. This was indeed the case with TF binding at regions within the *memory 2 and 3* clusters (Fig. 4A,4B, 4D). However, contrary to this expectation, our results reveal that AP-1 exhibits differential binding in activated memory T cells even at regions within the *memory 1* cluster lacking a poised chromatin state (Fig. 4A, 4B). Using a publicly available DNA methylation dataset from resting naive, TCM, and TEM CD4 T cells^10^ we assessed changes in CpG methylation associated with memory differentiation. We next examined CpG methylation dynamics within activation-induced accessible chromatin clusters and examined methylation patterns in our naive and memory cell types (Fig. 4A). Our analyses revealed a reduction in DNA methylation across the 3 memory clusters in memory T cells compared to naive T cells (Fig. 4A, 4C).

**Fig. 4:**
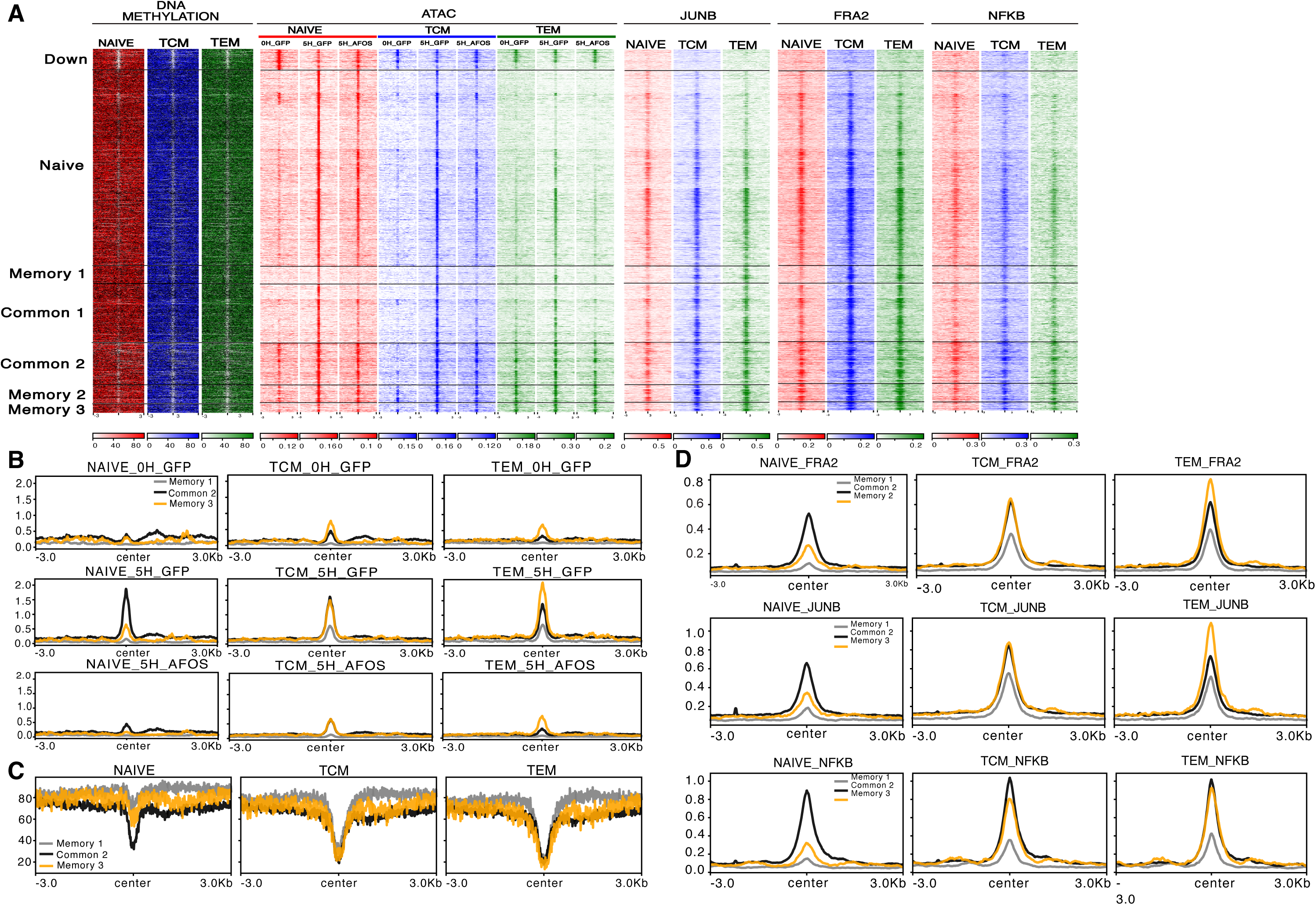
Reduced DNA methylation and accessible chromatin in memory T cells influence differential binding of AP-1 in memory T cells. (**A.**) Tag density heatmap showing the levels of DNA methylation and binding of JUNB, FRA2, NFAT, and NF-κB in regions with changed chromatin accessibility after 5-h activation in naive, TCM, and TEM cells. In resting memory cells, the *memory 2 and 3* clusters exhibit a poised chromatin state that was associated with increased AP-1 binding. In contrast, *memory 1* did not show chromatin posing at the level of chromatin accessibility. All three memory clusters show DNA demethylation in resting memory T cells. (**B-D.**) Profile plots showing three selected clusters: *memory 1*, *memory 3*, and *common 2* as a control in naive and memory cells. (**B.**) DNA accessibility. (**C.**) DNA methylation. (**D.**) JUNB, FRA2, and NF-κB binding.

We further assessed the effects of DNA methylation on the differential binding of AP-1 and NF κB in memory T cells. Specifically, we analyzed methylation patterns at naive-specific, memory-specific, and commonly bound AP-1 and NF κB sites across our naive, TEM, and TCM subsets. Our results revealed that memory-specific AP-1 (JUNB, FRA2) and NF κB binding sites exhibit markedly reduced DNA methylation in TEM and TCM cells compared to naive cells (Fig. 5A). In contrast, naive-specific and commonly bound AP-1 (FRA2, JUNB) and NF κB sites showed no significant differences in methylation between naive and memory T cell subsets (Fig. 4D). Collectively, these findings indicate that reduced DNA methylation and a poised epigenetic landscape in memory CD4 T cells facilitate differential TF binding, thereby contributing to memory-specific regulatory programs.

**Fig. 5:**
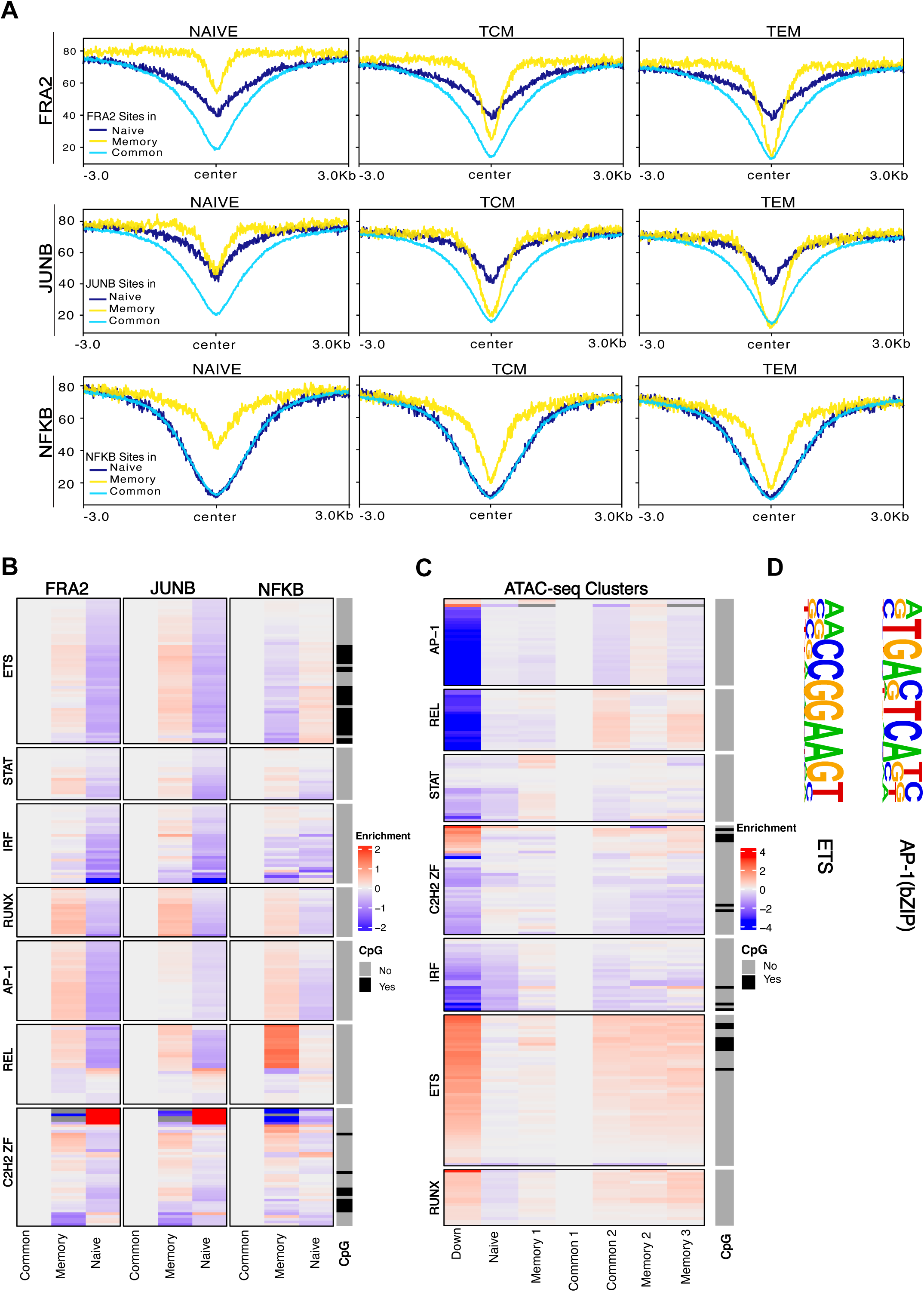
Memory-specific, AP-1‒bound sites have reduced DNA methylation and are enriched for AP-1 co-factors. (**A.**) Methylation profile plots showing the levels of DNA methylation around common, naive-specific, and memory-specific JUNB, FRA2, and NF-κB binding sites in naive TCM and TEM cells (**B-C.**) Heatmap showing motif enrichment within ATAC-seq peak clusters (**B**.) and ChIP-seq clusters (**C.**). Enrichment is normalized to the control peak set *common 1* for ATAC-seq clusters and the common peaks for ChIP-seq clusters. Presence of CpG dinucleotide within motifs is highlighted on the right. (**D.**) AP-1 and ETS transcription factor family motifs. Some ETS motifs have a CpG dinucleotide and can be affected by DNA methylation.

### AP-1 co-factors may contribute to differential AP-1 binding in memory T cells

DNA methylation is known to affect chromatin compaction of whole regions and accessibility of regulatory elements to TFs^29,30^ However, we wanted to address the possibility that DNA methylation may influence the binding of AP-1 directly. AP-1 DNA binding motifs lack CpG dinucleotides; therefore, AP-1 binding should not be directly affected by DNA methylation. However, AP-1 is known to bind DNA together with various cell type specific co-factors^31^. To elucidate the molecular factors underlying the selective binding of AP-1 in memory T cells, particularly at memory-specific regions with reduced CpG methylation, we sought to identify TFs that may function as AP-1 co-factors in memory T cells. To this end, we examined the distribution of AP-1 co-factor motifs across naive-specific, memory-specific, and common JUNB and FRA2-bound sites. This comparative motif enrichment analysis revealed that ETS, RUNX, STAT, nuclear receptor, and certain C2H2 ZF motifs were significantly enriched at memory-specific AP-1 binding sites relative to the commonly bound sites used as a control (Fig. 5B). We also performed TF motif enrichment analysis across all activation-induced chromatin accessible clusters compared to commonly accessible (c*ommon 1*) control regions to identify TFs enriched in naive and memory-specific clusters. Notably, memory-specific accessible chromatin clusters (*memory 1, 2, and 3*) were enriched for ETS, RUNX, STAT, and NF κB (REL) TF family motifs compared to the common clusters (Fig. 5C). In contrast, naive-specific accessible regions were enriched for motifs associated with IRF, homeodomain-containing factors, and nuclear factor families (Fig. 5C). Of interest, we observed enrichment of ETS motif variants that include a CpG dinucleotide (Fig. 5D), suggesting that binding of ETS factors to these regions may be directly affected by DNA methylation (Fig. 5D). Previous studies have reported a direct physical association of AP-1 and ETS TFs^32^ and have also shown that ETS factors are unable to bind CpG-methylated sites^33^. While the role of ETS factors will require further research, this suggests a mechanism wherein DNA methylation affects the binding of AP-1 by affecting the binding of its partners, possibly belonging to the ETS family. This is consistent with previous reports implicating ETS and RUNX TFs in rapid recall responses in memory T cells^34,35^. Collectively, these findings indicate that AP-1 binding in memory cells is shaped by both epigenetic features and TF co-factor binding, providing a possible mechanism for the epigenetic regulation of memory-specific immune response and rapid recall transcriptional programs.

### Memory-specific AP-1 binding sites overlap risk loci for immunologic diseases

Beyond their role in orchestrating protective immune response, CD4 T cells are essential regulators of immune homeostasis. Functional impairment of CD4 T cells can lead to autoimmune disorders, persistent infections, diminished immune responses, and malignancies^36^. We utilized regulatory element locus intersection (RELI) analysis^37^ to test whether the genetic variants located within naive-specific, common, and memory-specific JUNB, FRA2, and NF κB TF-bound sites and chromatin accessible regions are associated with susceptibility or protection against immune-related human diseases. Our analysis revealed a significant overlap between memory-specific and common chromatin accessible region, as well as binding sites of JUNB, FRA2, and NF κB, with genetic risk loci associated with multiple autoimmune and allergic diseases and other immune-related disorders (Fig. 6A, 6B). These include conditions such as multiple sclerosis, celiac disease, inflammatory bowel disease, and chronic inflammatory diseases (Fig. 6A, 6B).

**Fig. 6:**
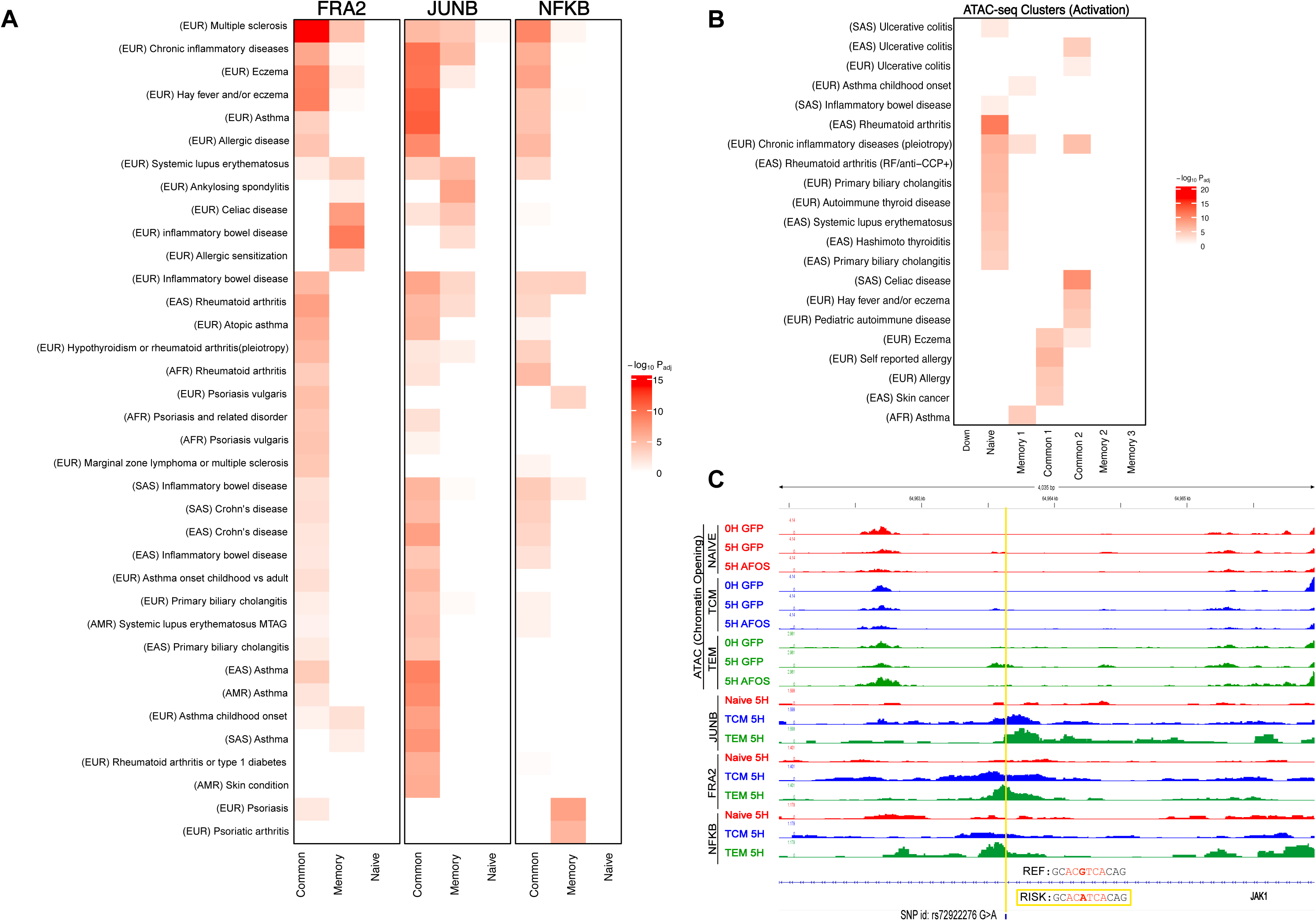
AP-1 binding sites in memory T cells coincide with genetic variants associated with immune-related diseases. (**A.**) Heatmap showing genetic disease association of cell-specific bound sites of JUNB, FRA2, and NF-κB. Values indicate the negative log of the corrected RELI p value (see Methods), which gauges the enrichment of genome-wide association study variants within each category. (**B.**) Heatmap showing disease association of ATAC-seq clusters and AP-1‒dependent accessible clusters. (**C.**) Genome browser tracks showing the genetic variant rs72922276, which is located in the *JAK1* locus and is associated with the multiple sclerosis and autoimmune thyroid disease locus. JUNB and FRA2 bind here in memory but not naive T cells, and this variant is predicted to alter the binding of AP-1. REF, reference sequence; RISK, risk variant sequence; SNP, single-nucleotide polymorphism.

We further searched for immune-disease risk variants harboring mutations that alter AP-1 binding sites and assessed variation in the occupancy of AP-1 naive and memory T cells at these loci along with additional evidence supporting their clinical relevance to the disease. One example is a single-nucleotide polymorphism (SNP) rs72922276 (G > A) located within the intergenic region of the *JAK1* gene that is associated with multiple sclerosis (MS, p = 1 x 10^-15^)^38^ and autoimmune thyroid disease (p = 5 x 10^-10^)^39^. Genome-wide association study (GWAS) population studies showed the presence of this SNP in European and African populations. A prior study also showed that the G:A mutation results in increased *JAK1* mRNA levels in CD8 and CD4 T cells^40^. Our data indicate binding of AP-1 in this region in memory T cells but not naive T cells (Fig. 6C). The SNP alters a putative AP-1 binding site and is predicted to abrogate AP-1 binding (Fig. 6C). Overall, our data show that memory-specific AP-1 binding may be important for immune system homeostasis.

## DISCUSSION

Previously, we showed that AP-1 is required for chromatin remodeling and the induction of gene expression upon naive CD4 T cell activation^18^. Here, we report that AP-1 is also required for the induction of gene expression in memory T cells. Among the AP-1 dependent genes are the genes involved in rapid recall in effector and central memory CD4 T cells. We found that enhanced inducibility of rapid recall genes in memory compared to naive cells is associated with differential binding of AP-1 to regulatory regions, including promoters and enhancers of rapid recall genes in TEM and TCM cell subsets compared to naive cells. Why is AP-1 able to bind at these new sites in memory but not in naive cells? At some of the new sites, this enhanced binding in memory T cells is consistent with the preestablished open chromatin state at the immune response loci likely retained from previous antigen encounters. However, at other sites, we observed no change in chromatin accessibility in resting memory compared to naive cells. Those sites, however, showed decreased CpG methylation in memory T cells compared to naïve cells.

Interestingly, the AP-1 motif itself does not contain CpG dinucleotide and therefore cannot be directly affected by DNA methylation. DNA methylation can affect the accessibility of the locus in general. However, in this case, our bioinformatics analysis points to a potential direct mechanism wherein DNA methylation affects binding of AP-1 binding partners, such as ETS family members. Indeed, AP-1 was previously shown to co-bind DNA together with multiple partners, including ETS, and RUNX, in T cells^31,32,41,42^. Our analysis showed enrichment of RUNX and ETS TF motifs in memory-specific AP-1 bound sites. The ETS motifs also contain CpG dinucleotides, suggesting reduced methylation at these CG sites may influence AP-1 binding in these memory-specific co-bound sites by affecting the binding of its ETS family partners. Of note, previous studies established that the DNA binding of the ETS family can be affected by DNA methylation^33^. Of course, establishing the role of ETS proteins in AP-1 binding will require further studies. Is the differential binding of AP-1 relevant to human health? We report that both common and memory-specific AP-1 binding sites are enriched for SNPs associated with human immune-related diseases, suggesting that proper functioning of these regulatory elements is important for immune homeostasis. Collectively, these findings show that the rapid recall ability of T cells is associated with differential binding of activation-inducible TFs, which is in turn modulated by the memory epigenome. This expands the understanding of AP-1 as a key regulator for immune response and rapid recall response in memory T cells.

## METHODS

### Isolation, culture, and activation of human CD4T cells

Peripheral blood mononuclear cells (PBMCs) were isolated from Lymphocyte Reduction Filters obtained from deidentified healthy human donors at the Hoxworth Blood Center of the University of Cincinnati. CD4 T cells were enriched by magnetic negative selection using EasySep Human CD4 Isolation Kit (Stemcell #17952) per manufacturer’s recommendations. To isolate specific subpopulations of CD4 T cells (Fig. 1B), enriched total CD4 T cells were stained with PE-Cy7 mouse anti-human CD45RO (#560608; BioLegend), FITC anti-human CD4 (#310904; BioLegend), and APC anti-human CD27 (#20-0279, Biolegend) antibodies. Flow sorting was performed on the BD FACS ARIAII, and analytical cytometry was performed on the BD FACS Canto II using the BD FACSDIVA to analyze the data. Purified CD4 T cells were cultured in Optimizer CTS (Gibco #A1048501) with 2 mM L-Glutamine supplement (Sigma # G7513) and 1X Penicillin-Streptomycin (Sigma # P4333) in incubators maintained at 37°C with 5% CO_2_. For T cell activation, prewashed human T cell activator anti-CD3/CD28 antibody beads (#11131D) were directly added to cultures of human CD4 T cells per manufacturer recommendation. Cultures were swirled every 10 min until the cells were bound to the beads (∼ 1 h) and incubated for 5 h.

### ChIP and library preparation by ChIPmentation

Fixing solution (11X: 50 mM Hepes-KOH, pH 8.0, 100 mM NaCl, 1 mM EDTA, 0.5 mM EGTA, and 8.8% formaldehyde) was added directly to T cell culture to a final formaldehyde concentration of 0.88% for fixation. After the incubation for 4 min on ice, 2.5M glycine solution was added to a final concentration of 125 mM and incubated at room temperature for 10 min to stop fixation. Cells were transferred to tubes and washed with cold PBS twice. All buffers used in the following procedures were supplemented with 1X protease inhibitor solution (#P8340; Sigma) before use. Fixed cells were resuspended in L1 buffer (50 mM Hepes-KOH, pH 8.0, 140 mM NaCl, 1 mM EDTA, 10% glycerol, 0.5% NP-40, 0.25% Triton X-100, and 1X protease inhibitors) and incubated on a rotator at 4°C for 10 min. Isolated nuclei were incubated with L2 buffer (10 mM Tris-HCl, pH 8.0, 200 mM NaCl, 1 mM EDTA, and 0.5 mM EGTA) with rotation for 10 min at room temperature. Then, isolated nuclei were carefully washed and resuspended in Tris-EDTA + 0.1% SDS solution. Cells were sonicated (peak power 105.0, duty factor 10.0, and cycle/burst 200) for 45 s in microtubes at 4°C using a S220 focused ultrasonicator (Covaris) to obtain the 200-bp to 500-bp fragments of chromatin. The chromatin solution was centrifuged, and the supernatant chromatin was collected. Triton X-100, glycerol, NaCl, and sodium deoxycholate were added into the chromatin solution to final concentrations of 1%, 5%, 150 mM, and 0.1%, respectively. Protease inhibitors (#P8340; Sigma) were also added.

ChIPs were performed in an IP-Star Compact automation system (Diagenode). In brief, 5–10 µg chromatin solution and 5–10 µL Protein A or G Dynabeads (Thermo Fisher Scientific) were used per reaction. Antibodies against FRA2 (Cell Signaling, D2F1E), JunB (Cell Signaling C37F9), and NF κB (Cell Signaling #3035) were used. The Dynabeads were sequentially washed with wash buffer 1 (RIPA 150 mM NaCl: 10 mM Tris-HCl, pH 8.0, 150 mM NaCl, 1 mM EDTA, 0.1% SDS, 0.1% sodium deoxycholate, and 1% Triton X-100), wash buffer 2 (RIPA 250 mM NaCl), wash buffer 3 (50 mM Tris-HCl, pH 8.0, 2 mM EDTA, and 0.2% N-Lauroylsarcosine sodium salt), and wash buffer 4 (TE + 0.2% Triton X-100) for 15 min each. Washed Dynabeads were processed with transposase per ChIPmentation protocol to accomplish tagmentation of ChIPed DNA^9^. Tagmented DNA was incubated with proteinase K in elution buffer (TE with 250 mM NaCl and 0.3% SDS) for 4 h at 65°C and purified from beads with the Qiagen MinElute DNA kit. PCR amplification and purification were performed in the same way as for ATAC-seq.

### ATAC-seq

Fifty thousand CD4 T cells were collected and processed in a transposase reaction, and library preparation was performed as described in the Omni-ATAC protocol^43^.

### GFP and A-FOS protein purification

Proteins were purified as described earlier^18^. A-FOS and GFP cDNA fragments were amplified from CMV500 A-FOS (#33353; Addgene) and pmaxGFP (Lonza) vectors by PCR, and a 6xHis-GST DNA fragment was also amplified from the pGEX vector. A-FOS, GFP, and 6xHis-GST were individually mixed with digested plasmid DNA (GE Healthcare Life Sciences) with *Nco*I restriction enzyme in an InFusion cloning reaction (TakaraBio). The resulting plasmid was transformed into the BL21 bacterial strain. A single colony was picked and transferred to 10 mL TB-SB (Terrific Broth–Super Broth) medium (TB/SB 1:1) with kanamycin, cultured overnight at 37°C, transferred to 500 mL TB-SB medium, and cultured until the optical density at 600 nm reached 0.8. For protein expression, isopropyl β-D-1-thiogalactopyranoside was added into the culture to a final concentration of 0.5 mM, and bacteria were cultured for another 6–8 h at 34°C. The bacteria were spun down, washed with PBS twice, and stored at -80°C. The bacterial pellet was resuspended in 20 mL lysis buffer (50 mM Na2HPO4, pH 8.0, 300 mM NaCl, 10% glycerol, 0.1% Triton X-100, and 20 mM imidazole with the addition of protease inhibitors [#04693159001; Roche], lysozyme [100 µg/mL], DNaseI [0.2 μg/mL], and β-mercaptoethanol to 1 mM), sonicated with a probe sonicator, and then frozen, thawed, and sonicated again.

The lysate was cleared by centrifugation and loaded on a 5-mL HisTrap HP column (#17524801; GE Healthcare Life Sciences) in Buffer A (50 mM Na2HPO4, pH 8.0, 300 mM NaCl, and 20 mM imidazole) using AKTA Start (GE Healthcare Life Sciences). The bound proteins were eluted by the gradient of Buffer B (50 mM Na2HPO4, pH 8.0, 300 mM NaCl, and 500 mM imidazole). The fractions containing protein were combined and subjected to dialysis against PBS with 1 mM β-mercaptoethanol overnight. After dialysis, Triton X-100 was added to the protein solution, and the His6-GST tag was cut with His-tagged HRV3C protease (#SAE0045; Sigma) at room temperature for 5 h. To remove the tags and the protease, imidazole was added to 20 mM, the solution was again passed through a 5-mL HisTrap HP column, and the flowthrough was collected. The purity was confirmed by SDS-PAGE (data not shown). The Endotoxin Removal Beads (#130-093- 657; Miltenyi Biotec) were used to remove endotoxins.

### GFP and A-FOS electroporation

A-FOS and GFP proteins were electroporated using Neon Nucleofector (Invitrogen) and the Neon Transfection System 10 µL Kit (#MPK1025; Invitrogen). Isolated naive, TEM, and TCM CD4 T cells (350,000 cells) were mixed with 2.5 µg A-FOS or GFP in T cell buffer and pulsed (voltage 2,400 V, width 20 ms, one pulse). For activation, the cells were stimulated 2 h after electroporation as described above.

### RNA isolation and RNA sequencing (RNA-seq)

Total RNA was isolated using the Monarch Total RNA Miniprep Kit (#T2010S), including on-column DNaseI digest. PolyA selection and fragmentation of RNA was conducted according to the NEBNext Ultra II RNA library prep kit (#E7775S). The fragmented RNA was subjected to first-strand and second-strand cDNA synthesis and adapter ligation according to the NEBNext Ultra II RNA library preparation for Illumina protocol.

### Sequencing data analysis

#### RNA-seq data preprocessing, alignment, and gene expression quantification

For each of the 27 samples, raw RNA-seq data in the form of paired-end FASTQ files were processed using “Trim Galore RNA-seq pipeline paired-end” (https://github.com/Barski-lab/workflows-datirium/blob/master/workflows/trim-rnaseq-pe.cwl) with default parameters (reporting only uniquely mapped reads with a maximum of 5 mismatches) in the SciDAP (https://scidap.com/) data analysis platform. Reads were trimmed with Trim Galore^44^ to remove adapter sequences and then aligned to the reference genome (GRCh38/hg38, https://www.ncbi.nlm.nih.gov/datasets/genome/GCF_000001405.26/) with STAR^45^. Gene expression levels were quantified using refGene annotations (https://hgdownload.cse.ucsc.edu/goldenPath/hg38/database/refGene.txt.gz).

#### RNA-seq differential expression analysis

The initial step for differential expression analysis was performed in SciDAP using the “DESeq2 LRT Step 1” pipeline (https://github.com/Barski-lab/workflows-datirium/blob/master/workflows/deseq-lrt-step-1.cwl). Gene expression from all 27 RNA-seq samples were processed together, modeling both additive and synergetic effects between the cell types (naive, TCM, and TEM) and the activation conditions (0 h GFP, 5 h GFP, and 5 h A-FOS). Genes with expression levels below 3 RPKM across all samples were excluded from the analysis. Additionally, the batch effect, defined by donor, was modeled in DESeq2 and then corrected using the Limma R package^46^ on heatmaps only. Targeted analysis for the specific pairwise contrasts (5 h GFP vs. 0 h GFP for each cell type) was conducted using the “DESeq2 Step 2 Wald” pipeline (https://github.com/Barski-lab/workflows-datirium/blob/master/workflows/deseq-lrt-step-2.cwl). All three target contrasts were processed in a single workflow run. Significant differentially expressed genes were selected using the following cutoffs for each contrast: adjusted p ≤ 0.1 and absolute log_2_ FC ≥ 0.585. The final results included only differentially expressed genes that passed filtering in at least one of the target contrasts.

#### Heatmap generation

Count data were vst-transformed in DESeq2 and minimum–maximum scaled across genes. Donor was included as a covariate in the DESeq2 model; prior to heatmap plotting, batch-corrected expression values were obtained using Limma’s removeBatchEffect() on the normalized counts. Hierarchical clustering was performed with HOPACH (Euclidean distance, average linkage, k = 5, kmax = 3), and heatmaps were generated using the pheatmap R package (v1.0).

#### ChIP-seq data preprocessing, alignment, and peak calling

For each of the 27 samples, raw ChIP-seq data in the form of paired-end FASTQ files were processed using the “Trim Galore ChIP-seq pipeline paired-end” (https://github.com/Barski-lab/workflows-datirium/blob/master/workflows/trim-chipseq-pe.cwl) with default parameters (reporting only uniquely mapped reads with a maximum of 3 mismatches) in the SciDAP (https://scidap.com/) data analysis platform. Reads were trimmed with Trim Galore^44^ to remove adapter sequences and then aligned to the reference genome (GRCh38/hg38, https://www.ncbi.nlm.nih.gov/datasets/genome/GCF_000001405.26/) with Bowtie^47^. PCR optical duplicates were removed with SAMtools^48^. The remaining reads were used for narrow peak calling with MACS2^49^. Only peaks with a q ≤ 0.05 were reported. The nearest genes were assigned using refGene annotations (https://hgdownload.cse.ucsc.edu/goldenPath/hg38/database/refGene.txt.gz).

#### ChIP-seq differential binding analysis

The initial step for differential binding analysis was performed in SciDAP using the “DiffBind LRT Step 1” pipeline (https://github.com/Barski-lab/workflows-datirium/blob/master/workflows/diffbind-lrt-step-1.cwl). Samples from each TF group (9 x FRA2, 9 x JUNB, and 9 x NF κB) were analyzed independently, modeling only the additive effect between the cell types (naive, TCM, and TEM) and donors. Peaks loaded from the samples were first filtered by q ≤ 10^-5^ and then overlapped within each cell type, retaining only those reproducible in at least two samples. Targeted analyses for the specific pairwise contrasts (TCM vs. naive, TEM vs. naive, and TEM vs. TCM) were conducted using the “DiffBind Step 2 Wald” pipeline (https://github.com/Barski-lab/workflows-datirium/blob/master/workflows/diffbind-lrt-step-2.cwl). For each TF group, all three target contrasts were processed in a single workflow run. Significant differentially bound regions were selected using the following cutoffs for each contrast: adjusted p ≤ 0.1 and absolute log_2_ FC ≥ 0.585. The final results included only differentially bound regions that passed filtering in at least one of the target contrasts.

#### ATAC-seq data preprocessing, alignment, and peak calling

For each of the 27 samples, raw ATAC-seq data in the form of paired-end FASTQ files was processed using “Trim Galore ATAC-seq pipeline paired-end” (https://github.com/Barski-lab/workflows-datirium/blob/master/workflows/trim-atacseq-pe.cwl) with default parameters (reporting only uniquely mapped reads with a maximum of 3 mismatches) in the SciDAP (https://scidap.com/) data analysis platform. Reads were trimmed with Trim Galore^44^ to remove adapter sequences and then aligned to the reference genome (GRCh38/hg38, https://www.ncbi.nlm.nih.gov/datasets/genome/GCF_000001405.26/) with Bowtie^47^. PCR optical duplicates were removed with SAMtools^48^. The remaining reads were used for narrow peak calling with MACS2^49^. Only peaks with a q ≤ 0.05 were reported. The nearest genes were assigned using refGene annotations (https://hgdownload.cse.ucsc.edu/goldenPath/hg38/database/refGene.txt.gz).

#### ATAC-seq differential accessibility analysis

The initial step for differential accessibility analysis was performed in SciDAP using the “DiffBind LRT Step 1” pipeline (https://github.com/Barski-lab/workflows-datirium/blob/master/workflows/diffbind-lrt-step-1.cwl). Chromatin accessibility data (peaks and reads) from all 27 ATAC-seq samples were processed together, modeling both additive and synergetic effects between the cell types (naive, TCM, and TEM) and the activation conditions (0 h GFP, 5 h GFP, and 5 h A-FOS). Peaks loaded from the samples were first filtered by q ≤ 10^-5^ and then overlapped within each group defined by cell type and activation condition, retaining only those reproducible in at least two samples. Additionally, the batch effect, defined by donor, was corrected using the Limma R package^46^. Targeted analyses for the specific pairwise contrasts (5 h GFP vs. 0 h GFP for each cell type) were conducted using the “DiffBind LRT Step 2” pipeline (https://github.com/Barski-lab/workflows-datirium/blob/master/workflows/diffbind-lrt-step-2.cwl). All three target contrasts were processed in a single workflow run. Significant differentially accessible regions were selected using the following cutoffs for each contrast: adjusted p ≤ 0.1 and absolute log_2_ FC ≥ 0.585. The final results included only differentially accessible regions that passed filtering in at least one of the target contrasts.

#### Gene ontology analysis

Gene ontology analysis was conducted using ToppGene^50^ for figures 1, S1 and S2 (expression data) and using annotatePeaks.pl function in Homer for figures 3 and S6 (ATAC peaks).

#### Gene set enrichment analysis

Naive, common, and memory-specific regions from ATAC-seq and ChIP-seq were assigned to genes by linear proximity (± 10 kb from TSS) and 3D chromatin looping datasets^26^ and microC^27^ as summarized in Table S9^51^.These genes were used to create cluster-specific gene sets. Genes generated from normalized read counts from differential gene expression analysis between naive and TEM cells were imported into the gene set enrichment analysis (GSEA) tool^28^ and are utilized as ranked gene lists for GSEA.

#### Whole-genome bisulphite sequencing analysis

Previously published bisulphite (BS)-seq data^10^ (EGAS00001001624) were reanalyzed for this project in the following way: preprocessing, alignment, and cytosine methylation calling for each of the 6 samples of naive, TEM, and TCM raw BS-seq data files were performed using the “Bismark Methylation PE” pipeline (https://github.com/Barski-lab/workflows-datirium/blob/master/workflows/bismark-methylation-pe.cwl) with default parameters in the SciDAP (https://scidap.com/) data analysis platform. Reads were trimmed with Trim Galore^44^ to remove adapter sequences and then aligned to the reference genome (GRCh38/hg38, https://www.ncbi.nlm.nih.gov/datasets/genome/GCF_000001405.26/) with Bismark^52^. Mapped reads were used for cytosine methylation calls and report generation.

#### Transcription factor motif enrichment analysis

Enrichment analysis of human TF motifs available in CisBP^53^ (build 3.00) was performed using a version of HOMER^54^ (v4.9.1) that was modified to employ a log_2_ likelihood scoring system. Enrichment scores were calculated for each motif as the log_2_ ratio of motif occurrence in target peaks vs. background. Relative motif enrichment was calculated by subtracting the control enrichment score (“activation-3” for ATAC-seq, “common” for each ChIP-seq TF). The top motifs for visualization were determined by filtering results for significance (q < 1 x 10^-6^) and then selecting the top 10 motifs per family and sample using the negative log_10_ p value.

#### Phenotype association analysis

The significance of the overlap between peak groups and variants associated with GWAS-derived phenotypes was performed as previously described^55^ using an updated GWAS catalog. In brief, the Genome Wide Association Studies Catalog^56^ (https://www.ebi.ac.uk/gwas/ v1.0.3.1) was downloaded on 2025-02-18 and parsed to generate a custom, ancestry-specific phenotype catalog. Risk loci for downstream analyses were identified using PLINK^57,58^ (v1.90b) to select independent risk variants through linkage disequilibrium pruning (r^2^ < 0.2) followed by linkage disequilibrium (LD) expansion (r^2^ > 0.8). RELI analyses were performed as previously described^37^ using the variant lists. Multiple-testing correction was applied using the Bonferroni method^59^, and records with less than 3 overlaps were assigned an adjusted p value of 1. Phenotypes were significantly associated with a peak set if the adjusted p value were less than 1 x 10^-4^.

### Antibodies List

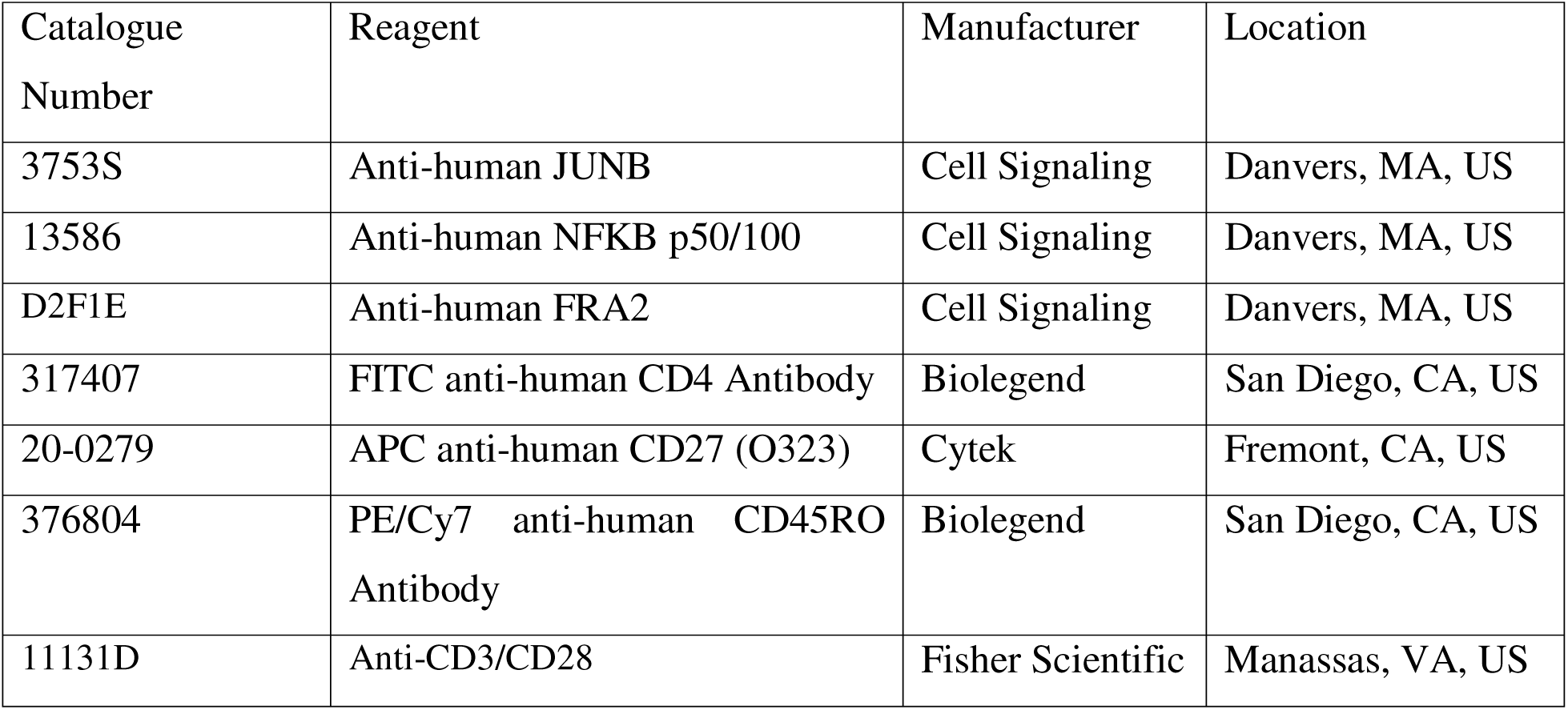

| Catalogue Number | Reagent | Manufacturer | Location |
| --- | --- | --- | --- |
| 3753S | Anti-human JUNB | Cell Signaling | Danvers, MA, US |
| 13586 | Anti-human NFkB p50/100 | Cell Signaling | Danvers, MA, US |
| D2F1E | Anti-human FRA2 | Cell Signaling | Danvers, MA, US |
| 317407 | FITC anti-human CD4 Antibody | Biolegend | San Diego, CA, US |
| 20-0279 | APC anti-human CD27 (O323) | Cytex | Fremont, CA, US |
| 376804 | PE/Cy7 anti-human CD45RO Antibody | Biolegend | San Diego, CA, US |
| 11131D | Anti-CD3/CD28 | Fisher Scientific | Manassas, VA, US |

### Reagents

| Catalogue Number | Reagent | Manufacturer | Location |
| --- | --- | --- | --- |
| 17952 | EasySep Human CD4 Isolation Kit | STEMCELL | Seattle, WA, USA |
| MT25072CV | Lymphocyte separation medium | Fisher Scientific | Manassas, VA, US |
| G7513 | L-Glutamine | Sigma-Aldrich | Burlington, MA, US |
| P4333 | Pennicilin & Streptomycin | Sigma-Aldrich | Burlington, MA, US |
| A1048501 | CTS Optimizer | Gibco | Waltham, MA, US |
| G7513 | L-Glutamine Supplement | Sigma-Aldrich | Burlington, MA, US |
| P4333 | Penicillin-Streptomycin Solution | Sigma-Aldrich | Burlington, MA, US |
| M0541S | NEBNext Ultra-II HiFidelity PCR Master Mix | New England Biolabs | Ipswich, MA, USA |
| M0544S | NEBNext Ultra-II Q5 PCR Master Mix | New England Biolabs | Ipswich, MA, USA |
| C01070012-30 | Tn5 Transposase Enzyme | Diagenode | Denville, NJ, USA |
| T2010S | Monarch Total RNA Miniprep Kit | New England Biolabs | Ipswich, MA, USA |
| E7775S | NEBNext Ultra II RNA Library Prep | New England Biolabs | Ipswich, MA, USA |
| T2010S | NEB Total RNA Miniprep Kit | New England Biolabs | Ipswich, MA, USA |
| 10002D | Dynabeads Protein A | Invitrogen | Vilnius, Lithuania |
| P8340 | Protease Inhibitor | Sigma | St. Louis, MO, USA |
| M0541S | High-Fidelity 2X PCR Master Mix | New England Biolabs | Ipswich, MA, USA |
| 28206 | MinElute Reaction Cleanup kit (250) | Qiagen | Hilden, Germany |
| 252549 | Formaldehyde (37%) | Sigma-Aldrich | Burlington, MA, US |
| A63880 | Ampure Beads | Beckman Coulter | Brea, CA, US |
| MPK 1025 | Neon Electroporation Kit, 100 uL Tips | Invitrogen | Waltham, MA, USA |
| 130-093- 657 | Endotoxin Removal Beads | Miltenyi Biotec | Gaithersburg, MD, US |
| 17524801 | HisTrap HP column | GE Healthcare Life Sciences | Boston, MA, US |
| SAE0045 | HRV3C protease | Sigma-Aldrich | Burlington, MA, US |

## Acknowledgements

We thank Masashi Yukawa, PhD for the purifying of the A-FOS and GFP proteins used for this project. We thank the University of Cincinnati Hoxworth Blood Center for providing of human blood samples, the CCRF Research Flow Cytometry Core (supported by National Institutes of Health [NIH] AR070549, NIDDK P30 DK078392); the University of Cincinnati Genomics, Epigenomics and Sequencing Core; and members of the Division of Allergy and Immunology at Cincinnati Children’s Hospital Medical Center for valuable discussions and technical and editorial assistance. This work was supported, in part, by the NIH grants R01 AI153442 and R42 HG011219 (A.B.).

## Ethics declarations

Cincinnati Children’s Hospital Medical Center Institutional Review Board approved the collection of deidentified human blood samples that was not deemed to be human subject research.

## Author information

### Materials and correspondence

Correspondence and material requests should be addressed to Artem Barski.

### Author affiliations

Division of Allergy and Immunology, Cincinnati Children’s Hospital Medical Center

Adenike R. Shittu, Michael Kotliar, Valerii Pavlov, Sarah Potter, Mathew T. Weirauch, Artem Barski

Development, Stem Cell and Regenerative Medicine Graduate Program, University of Cincinnati Adenike R. Shittu

Center for Autoimmune Genomics Etiology, Cincinnati Children’s Hospital Medical Center Andrew VonHandorf, Xiaoting Chen, Mathew T. Weirauch

Division of Human Genetics, Cincinnati Children’s Hospital Medical Center Mathew T. Weirauch, Artem Barski

Divisions of Biomedical Informatics and Developmental Biology, Cincinnati Children’s Hospital Medical Center

Mathew T. Weirauch

Department of Pediatrics, University of Cincinnati College of Medicine

Mathew T. Weirauch, Artem Barski

### Author contributions

A.B. and A.R.S. conceived and designed the experiments. A.R.S. conducted experiments with help from S.P and AB.; A.R.S, A.V., V.P., M.K., X.C., M.T.W., and A.B. conducted the analyses. A.R.S. and A.B. wrote the manuscript. A.R.S., A.V., S.P., V.P., M.K., X.C., M.T.W., and A.B. revised the manuscript. All authors approved the final version of the manuscript.

