## Supplemental figures for "Memory T cell rapid recall is driven by memory-specific AP-1 recruitment determined by epigenome and co-factor interactions"

**Supplementary Fig. S1**

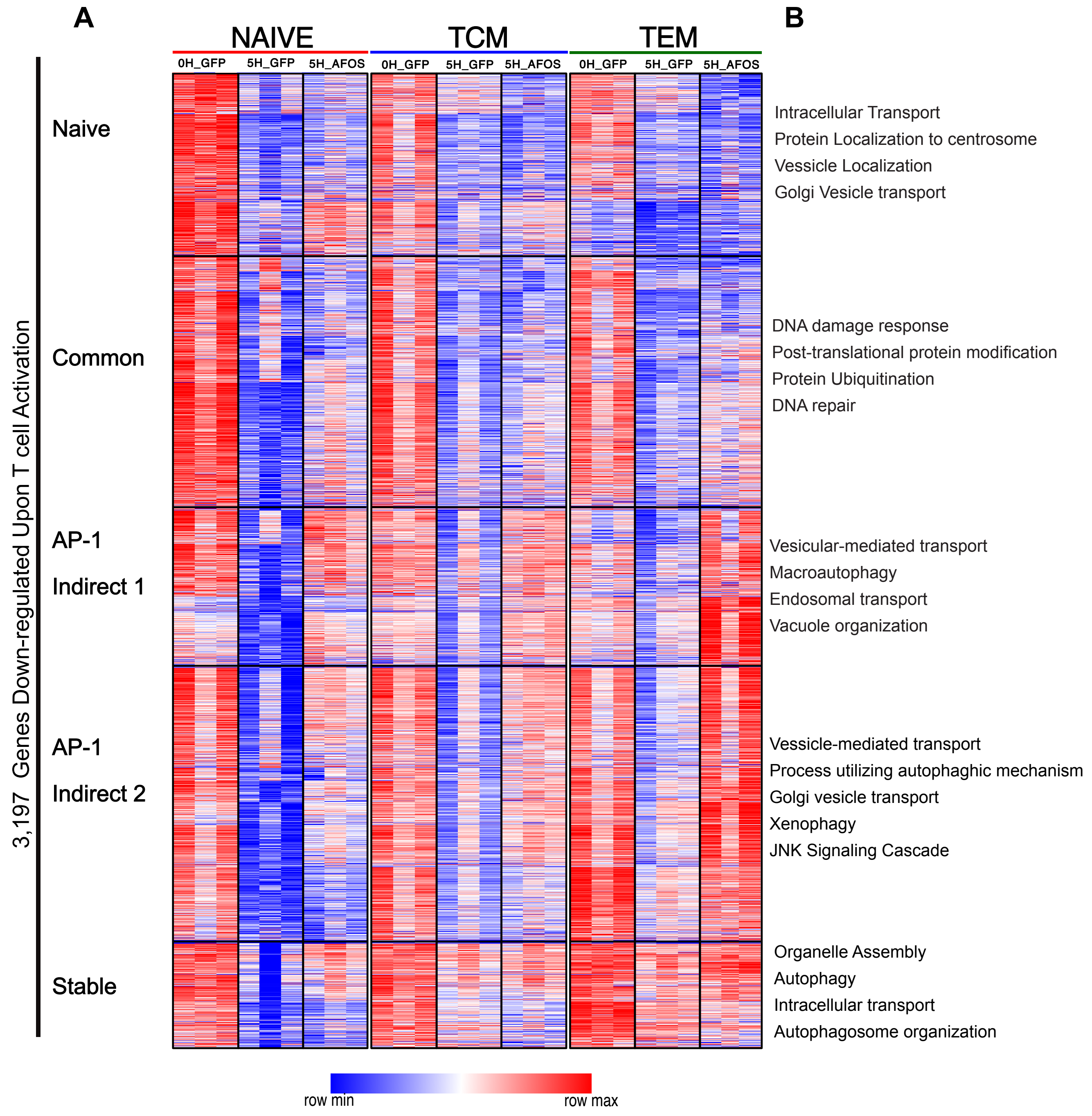

**Supplementary Fig. S1. Genes downregulated during activation show subset-specific expression differences in naïve versus memory T cells.** (**A.**) Heatmap depicting mRNA expression levels of activation-repressed genes in naïve, effector (TEM), and central (TCM) memory CD4 T cells. Gene sets are clustered using expression patterns across naïve and memory subsets. (B**.**) Pathway enrichment analysis of clusters of genes that are downregulated upon activation.

**Supplementary Fig. S2**

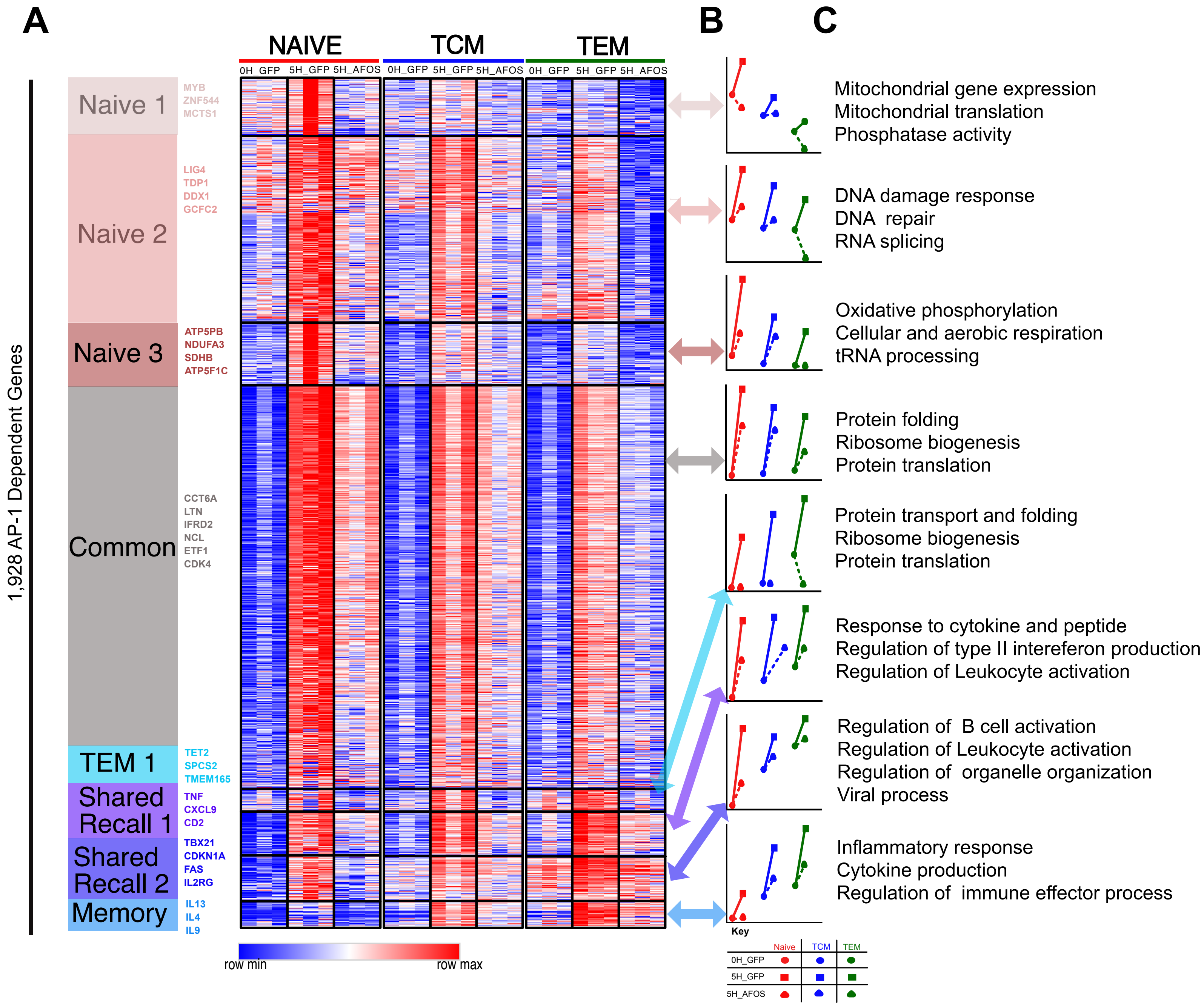

**Supplementary Fig. S2. AP-1–dependent differentially expressed genes in naïve and memory T cells.** (**A.**) Heatmap depicting mRNA expression levels of genes that are dependent on AP-1 in naïve, effector (TEM), or central (TCM) memory CD4 T cells. Gene sets are clustered using expression patterns across naïve and memory subsets. (**B.**) Line plot showing Z score–scaled average expression values for each cluster across cell types and conditions. (**C.**) Pathway enrichment analysis of AP-1–dependent gene clusters.

**Supplementary Fig. S3**

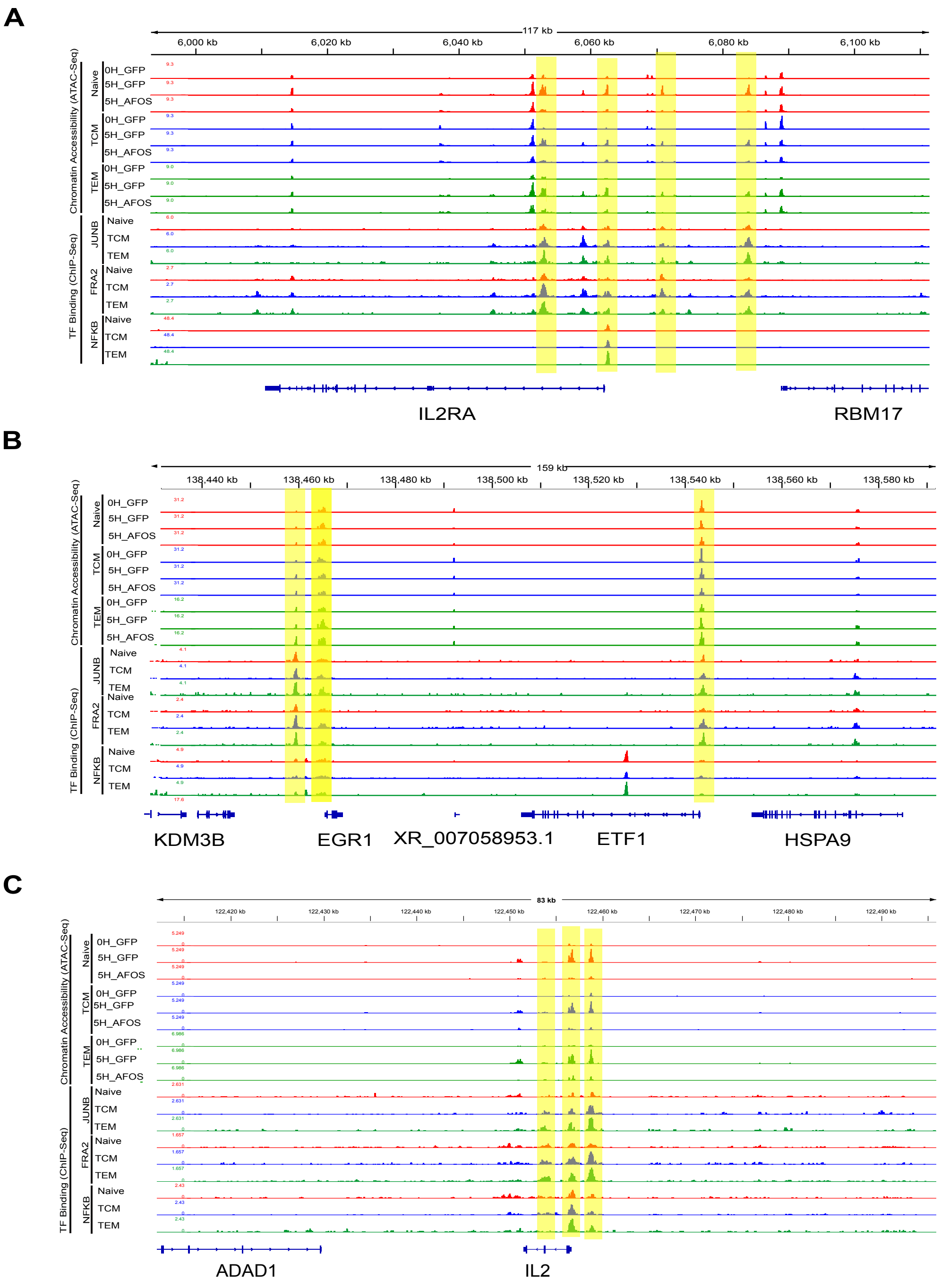

**Supplementary Fig. S3. Differential binding of AP-1 (JUNB and FRA2) and NF-κB in naïve and memory T cells.** (**A-B.**) Genome browser tracks showing chromatin accessibility (ATAC-seq) and AP-1 (JUNB and FRA2) and NF-κB binding at the *IL2RA*, *ETF1*, *EGR1*, and *IL2* locus in naïve, TEM, and TCM cells. TF, transcription factor. Regions of interest are highlighted

**Supplementary Fig. S4**

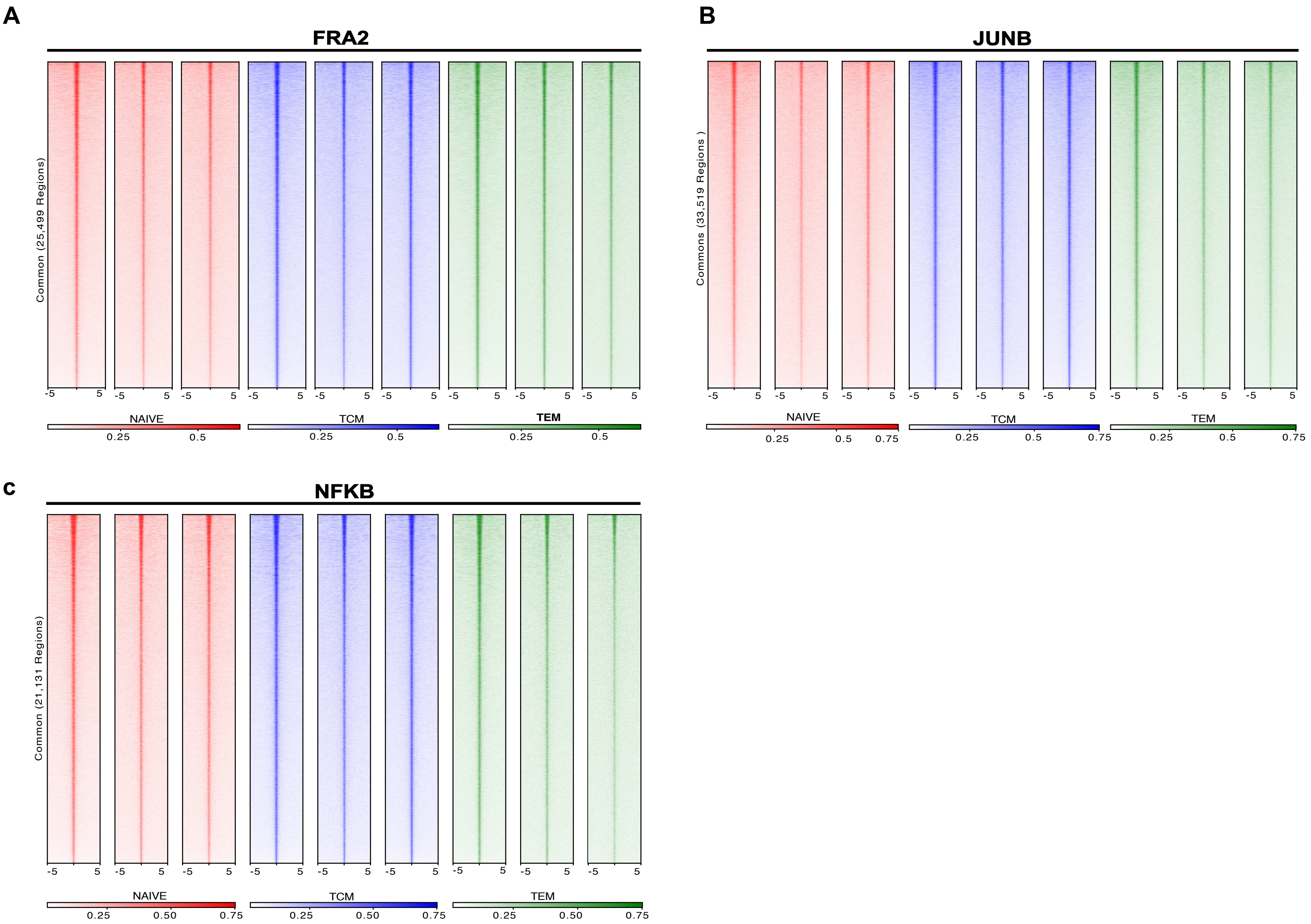

**Supplementary Fig. S4. Common AP-1– and NF-κB–bound sites in naïve and memory T cells.** (**A-C**.) Tag density plot illustrating the common bound sites of AP-1 (JUNB and FRA2) and NF-κB across naïve, TEM, and TCM subsets.

### Supplementary Fig. S5

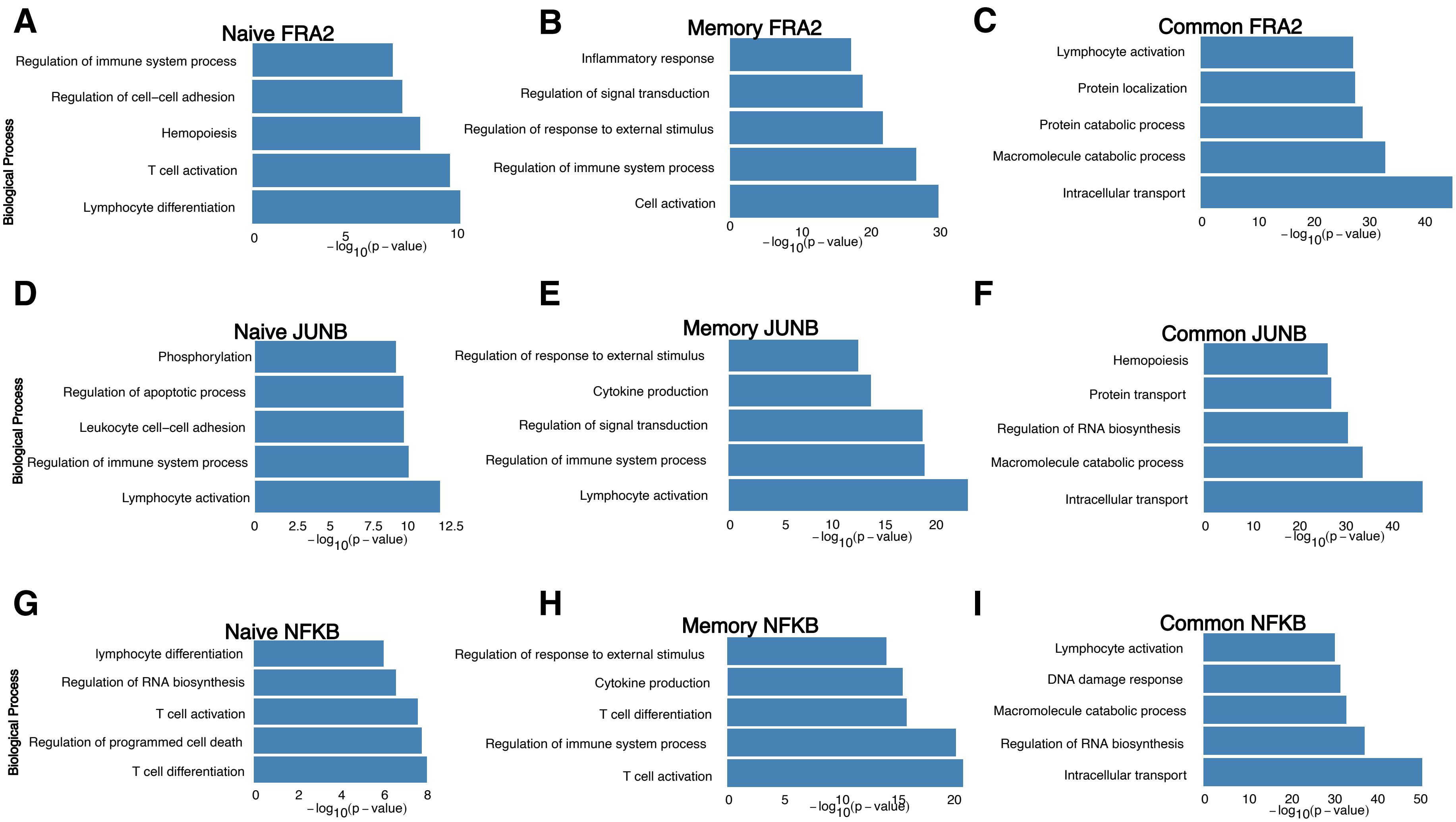

**Supplementary Fig. S5. Gene ontology analysis of naïve-specific, memory-specific, and common sites of AP-1 and NF-κB binding.** (**A-I**.) Bar plot illustrating the gene ontology of naïve-specific, memory-specific, and common bound sites of AP-1 (JUNB and FRA2) and NF-κB across naïve, TEM, and TCM subsets. P values generated are shown by ToppGene.

**Supplementary Fig. S6**

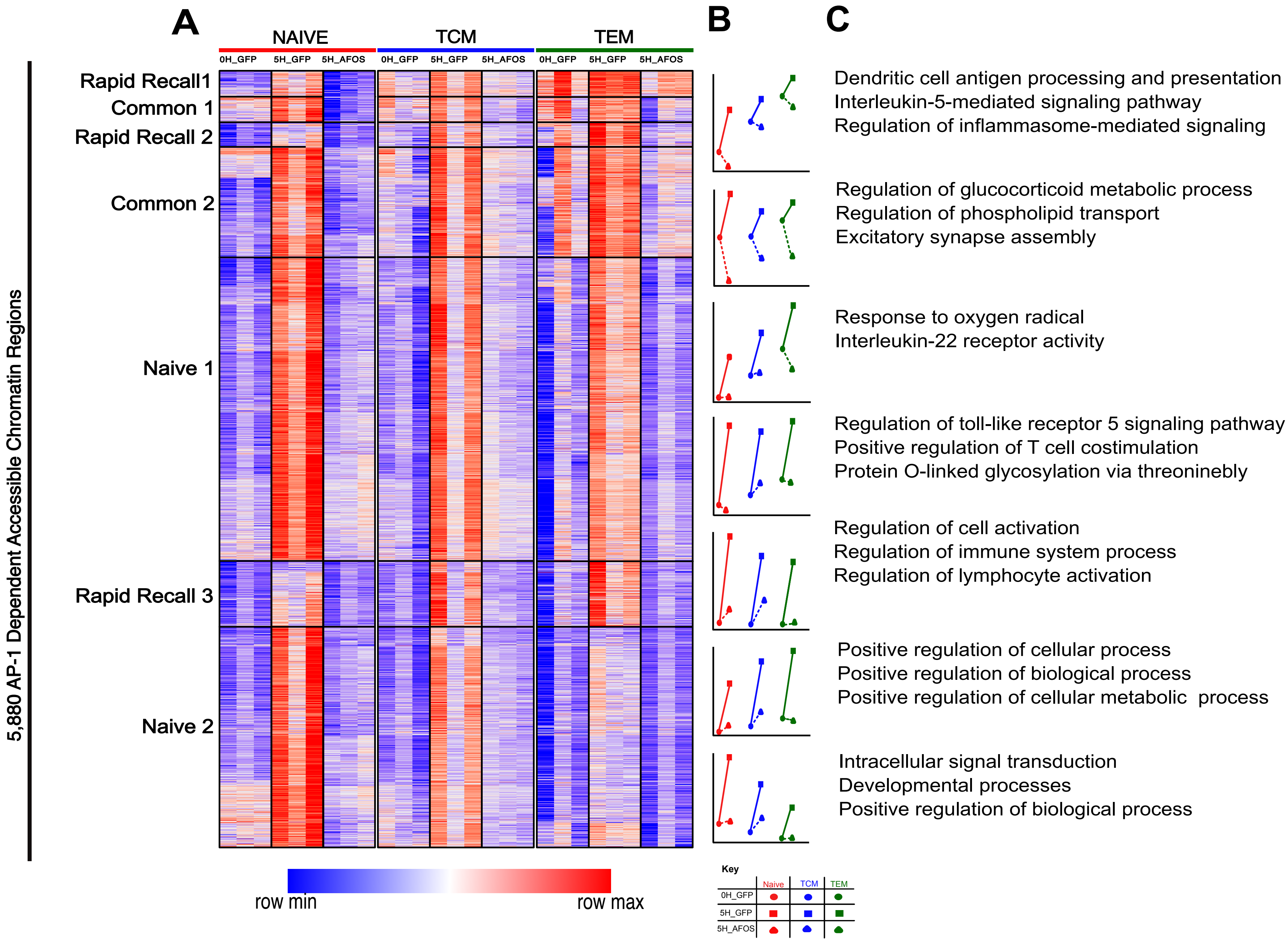

**Supplementary Fig. S6. AP-1–dependent chromatin remodeling in memory T cells.** (**A.**) Heatmap depicting chromatin-accessible regions that are dependent on AP-1 in naïve, effector (TEM), or central (TCM) memory CD4 T cells. Genomic regions are clustered by accessibility across naïve and memory subsets. (**B.**) Line plot showing Z score–scaled average expression values for each cluster across cell types and conditions. (**C.**) Pathway enrichment analysis of AP-1–dependent gene clusters.
